# Metacognitive awareness of difficulty in action selection: the role of the cingulo-opercular network

**DOI:** 10.1101/641340

**Authors:** Kobe Desender, Martyn Teuchies, Carlos Gonzalez Garcia, Wouter de Baene, Jelle Demanet, Marcel Brass

## Abstract

The question whether and how we are able to monitor our own cognitive states (metacognition) has been a matter of debate for decades. Do we have direct access to our cognitive processes or can we only infer them indirectly based on their consequences? In the current study, we wanted to investigate the brain circuits that underlie the metacognitive experience of fluency in action selection. To manipulate action-selection fluency we used a subliminal response priming paradigm. On each trial, both male and female human participants additionally engaged in the metacognitive process of rating how hard they felt it was to respond to the target stimulus. Despite having no conscious awareness of the prime, results showed that participants rated incompatible trials (during which subliminal primes interfered with the required response) to be more difficult than compatible trials (where primes facilitated the required response) reflecting metacognitive awareness of difficulty. This increased sense of subjective difficulty was mirrored by increased activity in the rostral cingulate zone (RCZ) and the anterior insula, two regions that are functionally closely connected. Importantly, this reflected activations that were unique to subjective difficulty ratings and were not explained by reaction times or prime-response compatibility. We interpret these findings in light of a possible grounding of the metacognitive judgement of fluency in action selection in interoceptive signals resulting from increased effort.

## 1. Introduction

To what extent are humans able to monitor their own cognitive processes? Looking at the literature there is some controversy surrounding this question. Some research suggests that humans are poor judges of their own cognitive processes (Nisbett & Wilson, 1977; Wilson & Dunn, 2004; Johansson, Hall, Sikström, & Olsson, 2005), whereas others have shown that humans are remarkably good at monitoring their own cognition, as participants are often aware when they made an error (Murphy, Robertson, Harty, & O’Connell, 2015), and can provide very precise estimates of the probability of being correct (Boldt & Yeung, 2015). This process of self-monitoring is also referred to as metacognition, as it describes insights into our own cognitive processes (Metcalfe & Shimamura, 1994; Brown, 1978). The neuro-cognitive mechanisms that underlie this self-monitoring ability are still poorly understood. Self-monitoring of behavior has been the focus of a number of recent brain imaging studies in the domain of error detection and decision making (Klein et al., 2007; Ullsperger, Harsay, Wessel, & Ridderinkhof, 2010; Fleming, Weil, Nagy, Dolan, & Rees, 2010). The anterior insula has been found to be involved in error detection (Klein et al., 2007; Ullsperger et al., 2010), whereas anterior prefrontal regions were found to be involved in decision confidence (Fleming et al., 2010). More recently, the process by which humans monitor the difficulty in action selection has attracted increasing attention (see e.g. Desender, Van Opstal, & Van den Bussche, 2014; Desender, Van Opstal, Hughes, & Van den Bussche, 2016; Questienne, Atas, Burle, Gevers, 2017). An interesting aspect of action selection is that one can manipulate its difficulty without participants becoming aware of the manipulation. It is known that subliminal response conflict hampers performance: it slows down response speed and it increases error rates (see e.g. Vorberg et al., 2003). By using a subliminal response priming paradigm, one can thus manipulate conflict between two response options outside participants’ awareness. Consequently, a metacognitive representation of this manipulation cannot be based on a conscious interpretation of the events (i.e., the visually conflicting information), but has to be based on the interpretation of internal signals caused by these events. A recent brain imaging study revealed that subliminal response conflict is registered in the brain by increased activity in both the rostral-cingulate zone (RCZ) and the left anterior insula (Teuchies et al. 2016). Even though participants are typically unaware of the subliminal conflict-inducing stimulus, previous research has shown that they nevertheless report increased levels of subjective difficulty when responding to trials with subliminal conflict (Desender et al., 2014; 2016). This raises the question about the source of such metacognitive judgements. A previous study indicates that the experience of subjective difficulty does not simply reflect a read-out of response speed (Desender et al., 2016), but rather seems to be based on motor conflict induced by the subliminal primes (Questienne, Atas, Burle, & Gevers, 2017). This leads to the prediction that the metacognitive judgement is related to brain processes that are involved in subliminal conflict processing itself, namely the RCZ and the anterior insula (Teuchies et al., 2016), independently from reaction times and prime-response compatibility.

## 2. Method Section

### 2.1 Participants

Participants in this study were 30 Dutch-speaking students from Ghent University (19 female, mean age = 23.77 years, SD = 3.20); each one reported to be healthy and with no history of neurological, pain, or circulatory disorders and normal or corrected-to-normal vision. One participant was removed due to excessive head motion. All participants gave written informed consent, and the study was approved by the Medical Ethical Review Board of the Ghent University hospital, in accordance with the declaration of Helsinki. All participants were right-handed, as assessed by the Edinburgh Inventory (Oldfield, 1971), and were compensated thirty Euro for their participation.

### 2.2 Stimuli

Stimulus presentation and response registration was done using Tscope software (Stevens, Lammertyn, Verbruggen, & Vandierendonck, 2006). In the scanner, the task was presented using a Brainlogics 200MR digital projector that uses digital light processing (DLP) running at a refresh rate of 60 Hz with a viewing distance of 120 cm. Using DLP it took 1 ms to deconstruct the image on the screen allowing our subliminal primes to be presented with great precision. The mean presentation time was 18.00 ms (*SD* = 0.24; range 15.91 – 18.91 ms). Three types of grey colored primes were used (Figure 1): left or right pointing arrows or a neutral prime (which consisted of overlapping left and right pointing arrows). The primes were followed by superimposed metacontrast masks of the same luminance. The metacontrast masks were embedded within target arrows that pointed left or right. Primes subtended visual angles of 0.8° × 1.86°, and the targets 1.09° × 3.47°. Prime and target stimuli could appear randomly above or below a fixation cross at a visual angle of 1.38°. The unpredictable location was included to enhance the masking effect (Vorberg et al., 2003). A circular rating scale was adapted from Kahnt, Heinzle, Park, & Haynes (2011). The x and y coordinates of the mouse response were converted into polar coordinates ranging from 0 degrees (easiest) to 360 degrees (most difficult). The thickness of the scale increased with difficulty. The easiest point on the scale was the tail of the circle; the most difficult point was the thickest point of the circle. The orientation of the scale was randomly chosen on each trial so that the starting point of the scale was unpredictable. This prevented participants from preparing a motor response before seeing the actual scale.

**Figure 1.**
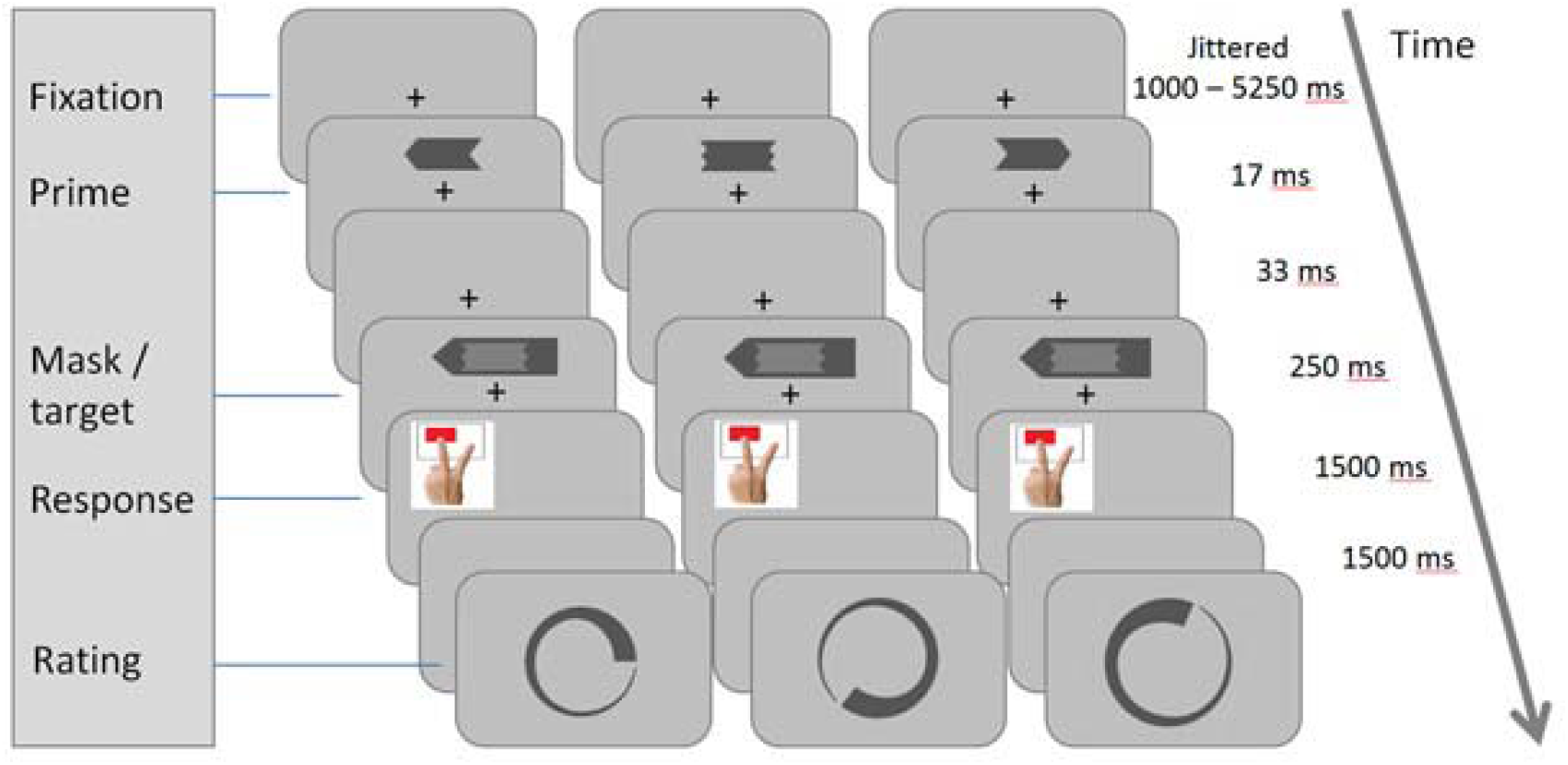
Schematic of an Experimental trial. Three possible combinations of the factor prime-response compatibility (compatible: left panel; neutral: middle panel; incompatible: right panel). Participants were instructed to respond to the target stimuli (with the left hand), and were unaware of the presence of the arrow primes. Primes and targets could appear randomly above or below fixation on each trial. After their response participants indicated their subjective feeling of difficulty using a circular rating scale. The thin tail is the easiest point and the scale continuously increases in thickness and difficulty up to the thick end representing the most difficult point. Participants were instructed to use the whole scale.

### 2.3 Procedure

Except for the ratings, the experimental design was identical to Teuchies et al. (2016). Primes were presented for 16.7 ms (1 refresh rate at 60Hz), followed by a blank screen for 33.3 ms and a target that also functioned as a mask. Target duration was 250 ms. The response window was set to 1500 ms. Participants were instructed to respond as fast and accurate as possible to the direction of the target arrows with their left middle finger (left pointing targets) and left index finger (right pointing targets) using an MR compatible response box. If participants failed to respond within this time window, they saw “te laat” (too late) for 1000 ms after the trial. After each response a blank screen was shown for 1500 ms followed by the rating part of the trial during which the rating scale was shown until participants had given their response with their right hand using an MR compatible optical track-ball mouse to select a point on the rating scale that matched their subjective sense of difficulty. The response was registered only when the mouse was actually on the rating scale. Mouse clicks outside of the rating scale were not registered. Participants were instructed to use the entire scale and they were informed that the extremities of the scale represented their personal most difficult and easiest points. Once they clicked on the scale a blank screen was shown for the inter-trial-interval. The inter-trial-interval was jittered with values ranging between 1000ms and 5250ms. The jitter values followed a distribution with pseudo-logarithmic density (range, 1000–5250 ms, in steps of 250 ms; mean jitter, 2625 ms).

Before doing the experiment in the scanner participants carried out two training blocks of 48 trials each. In the first training block they were only presented with the response priming task, without the rating to let them experience the response priming task. When asked, all participants indicated that they made mistakes and that some trials felt more difficult than others. In the second training block the rating was added after every individual trial and participants were instructed to rate on each trial how difficult they found it to respond as fast and accurate as possible to the target stimulus. Participants were never alerted to the possibility of primes being presented. The main task inside the MRI scanner consisted of three blocks of 72 trials each. Within each block, each prime-response compatibility condition (compatible, incompatible and neutral) occurred equally often. At the end of the task participants were asked whether they noticed anything unusual about the stimuli during the task. None of the participants noticed the primes, but three of them reported seeing a “flash” before the target was presented. These participants were included in the final sample. Following the test phase, participants were explicitly told about the presence of the primes and performed a prime-visibility test. This test allowed us to check if the prime stimuli were indeed presented subliminally, or not. In this test participants were asked to identify the direction of the primes (left or right) on each individual trial by using the same left and right response buttons as used during the test phase. During this test, participants remained in the scanner, so environment and apparatus were identical to the main experiment. To minimize indirect priming effects on the recognition of the primes, participants were required to respond at least 600 ms after the mask was presented. A visual cue (‘*’) signaled when they were allowed to respond. The test consisted of two blocks of 50 trials each.

### 2.4 fMRI data acquisition and preprocessing

Data were acquired with a 3T Siemens Magnetom Trio MRI system (Siemens Medical Systems, Erlangen, Germany) using a 32-channel radiofrequency head coil. Participants were positioned head first and supine in the magnet bore. First, 176 high-resolution anatomical images were acquired using a T1-weighted 3D MPRAGE sequence (TR = 2,250 ms, TE = 4.18 ms, TI = 900 ms, image matrix = 256 x 256, FOV = 256 mm, flip angle = 9°, and voxel size = 1 x 1 x 1 mm). Whole-brain functional images were then collected using a T2-weighted echo-planar imaging (EPI) sequence, sensitive to blood-oxygen-level dependent contrast (TR = 2,000 ms, TE = 35 ms, image matrix = 64 x 64, FOV = 224 mm, flip angle = 80°, slice thickness = 3.0 mm, distance factor = 17%, voxel size 3.5 x 3.5 x 3.0 mm, and 30 axial slices). A varying number of images were acquired per run due to individual differences in choice behavior and reaction times. All data were preprocessed and analyzed using Matlab and the SPM8 software (Wellcome Department of Cognitive Neurology, London, UK). To account for possible T1 relaxation effects, the first four scans of each EPI series were excluded from the analysis. The ArtRepair toolbox for SPM was used to detect outlier volumes concerning global intensity or large scan-to-scan movement (Mazaika, Whitfield-Gabrieli, & Reiss, 2007). First, a mean image for all scan volumes was created, to which individual volumes were spatially realigned using rigid body transformation. Thereafter, they were slice time corrected using the first slice as a reference. The structural image of each participant was co-registered with their mean functional image after which all functional images were normalized to the Montreal Neurological Institute (Montreal, Quebec, Canada) T1 template. Motion parameters were estimated for each session separately. The images were resampled into 3 x 3 x 3 mm voxels and spatially smoothed with a Gaussian kernel of 8 mm (full-width at half maximum). A high-pass filter of 128 Hz was applied during fMRI data analysis to remove low-frequency drifts.

### 2.5 Behavioral Data Analysis

Mean reaction times (RTs), error rates and subjective ratings were submitted to a repeated-measures ANOVA, with prime-response compatibility (compatible vs. incompatible vs. neutral) as factor. The responses to the primes in the visibility check were categorized using signal detection theory (Green & Swets, 1966). Measures of prime discriminability (*d’*) for each participant were computed. We then used a one-sample t-test to see whether the mean *d’* of the sample deviated from zero.

### 2.6 Regions of interest analyses

In the region of interest (ROI) analyses, we focused on the RCZ and the anterior insula as these were our principal ROI’s based on our previous study (Teuchies at al., 2016). Accordingly, the peak coordinates were taken from this previous study. To create ROI’s we created spheres with a 5mm radius around the peak coordinates of the RCZ [MNI 6 20 43] and the anterior insula [MNI -36 20 -2]. We then extracted single-trial beta estimates using a General Linear Model approach, in which each trial was modelled as one regressor. We used linear mixed models, as implemented in the lme4 package (Bates et al., 2015) to analyze the relationship between difficulty ratings and brain activity. Using this type of analysis, both variables can be fit at the single-trial level. Random slopes were added for each variable when this increased the model fit, as assessed by model comparison. For these models, *F* statistics are reported and the degrees of freedom were estimated by Satterthwaite’s approximation, as implemented in the lmerTest package (Kuznetsova et al., 2014).

### 2.7 GLM analyses

The participant-level statistical analyses were performed using the general linear model (GLM). In a first analysis, compatibility conditions (compatible, incompatible and neutral) were modelled in a single regressor of interest and raw subjective rating values for each trial were added as an extra parameter allowing us to look at brain activity related to the raw subjective difficulty ratings. In a second analysis, we wanted to capture variance in brain activity that was *unique* to the subjective difficulty ratings, independent of the variables prime-response compatibility and reaction time (which are both known to affect subjective difficulty ratings; Desender, Van Opstal, & Van den Bussche, 2017; Questienne, Atas, Burle, & Gevers, 2017). In order to capture variance related to compatibility, three different regressors of interest (compatible/incompatible/neutral) were modelled for this variable. In order to look at brain activation uniquely attributed to subjective ratings independent of reaction times (i.e., both variables showed modest negative relation: mean *r* = -.30, *sd* = .16, range = -.545 and .011), we introduced them both as parametric modulators. Because the order of the parametric regressors matters (i.e., the second regressor will only capture variance that has not been captured yet), we first entered reaction time as a parametric regressor and subjective rating as the second parametric regressor.

In both analyses, erroneous trials and the first trials of each block were always modeled as separate regressors of no interest (4.9% of the trials). The events of interest were the periods after the onsets of the different targets in the response priming task. Vectors containing the event onsets were convolved with the canonical hemodynamic response function (HRF) to form the main regressors in the design matrix (the regression model). Motion parameters for each individual subject were added. No derivatives were added to the model for this analysis. The statistical parameter estimates were computed separately for each voxel for all columns in the design matrix. Contrast images were constructed for each individual to compare the relevant parameter estimates for the regressors containing the canonical HRF. The group-level random effects analysis was then performed. Using one-sample t-tests we looked at the effects of the subjective difficulty ratings and reaction times across prime-response compatibility conditions. The subjective difficulty ratings and the reaction times had been added as parametric regressors during the first-level analysis. To correct for multiple comparisons, first we identified individual voxels that passed a ‘height’ threshold of p < 0.001, and then the minimum cluster size was set to the number of voxels corresponding to p < 0.05, FWE-corrected. This combination of thresholds has been shown to control appropriately for false-positives (Eklund et al., 2016). The resulting maps were overlaid onto a structural image of a standard MNI brain, and the coordinates reported correspond to the MNI coordinate system.

### 2.8 Mediation analyses

As described below, activity in both the RCZ and the anterior insula was related to difficulty ratings. To shed light on the direction of these effects, we performed post-hoc mediation analyses. For this analysis we created a new general linear model in which all the trials were entered as separate regressors, so we obtained brain activation for the RCZ and the anterior insula on a trial-by-trial basis. Our main question was whether the influence of the RCZ on difficulty ratings was mediated by the anterior insula, conditional on congruency. A mediator and an outcome model were fitted on the data using mixed regression modelling, using the same model building strategy as reported above. A mediator mixed model was fit in which activity in the anterior insula was predicted by activity in the RCZ, reaction times and compatibility. An outcome mixed model was fit in which difficulty ratings were predicted by activity in the anterior insula, the RCZ, reaction times and compatibility. A mediation analysis was then performed on these two models (using the mediation package; Tingley et al., 2014). This method partitions the total effect on ratings into an indirect effect (i.e. the effect of RCZ on ratings that is mediated by anterior insula) and a direct effect (i.e., correlation between RCZ and ratings that is not explained by anterior insula), conditional on reaction times and compatibility. If this indirect effect is significant, this is evidence for a significant mediation effect. Note that this latter observation is equivalent to showing that the influence of a direct path decreases when a mediation path is added to the model. Second, we tested the reversed hypothesis that the influence of anterior insula on difficulty ratings was mediated by the anterior insula. For this, the same mediation analysis was run but after exchanging RCZ and anterior insula in all models.

## 3. Results

### 3.1 Behavioral Results

#### 3.1.1 Main Task

Trials where participants did not respond within the 1500 ms response window were removed from the data (0.6% of the trials). For the remaining data, mean RTs on correct trials, mean error rates and mean difficulty judgments on correct trials were submitted to separate repeated-measures ANOVA’s with prime-response compatibility (prime-response compatible vs. incompatible vs. neutral) as factor. For RTs (table 1), this analysis yielded a significant effect of prime-response compatibility, (*F*(2, 28) = 39.24, *p* < .001, ηp^2^ = .737). Prime-compatible responses (*M* = 426.8 ms) were significantly faster than prime-incompatible responses (*M* = 453.9 ms; incompatible – compatible = 27 ms; *t*(29) = 7.94, *p* < .001, *d* = 0.68). Prime-compatible responses were not faster than prime-neutral responses (*M* = 430.6 ms; neutral – compatible= 7ms; *t*(29) = -1.27, *p* = .22, d = 0.11), meaning that directional primes did not lead to a significant facilitation effect. There was, however, a significant interference effect, meaning that prime-incompatible responses were slower than responses to neutral primes (incompatible – neutral = 23ms; *t*(29) = 7.97, *p* < .001, *d* = 0.62).

**Table 1.** Reaction times, percentage of errors and difficulty ratings as a function of prime-action compatibility.

|  | Reaction time (ms) | Errors (%) | Subjective difficulty rating (°) |
| --- | --- | --- | --- |
| Compatible | 426.8 (8.1) | 2.92 (0.6) | 146.3 (9.5) |
| Incompatible | 453.9 (6.8) | 7.93 (1.3) | 156.6 (9.5) |
| Neutral | 430.6 (6.7) | 3.84 (0.6) | 145.8 (9.4) |

Note: numbers in parentheses show standard errors of the means across participants. The subjective difficulty ratings are reported in degrees ranging from 0 – 359.

The error rates showed a similar effect of prime-response compatibility, (*F*(2, 28) = 12.53, *p* < .001, ηp^2^ = .472). Participants made significantly more errors on prime-incompatible trials (*M* = 7.93%) than on prime-compatible trials (*M* = 2.92%; *t*(29) = 5.1, *p* < 0.001, *d* = 0.93) and on neutral trials (*M* = 3.84%; *t*(29) = 4.3, *p* < 0.001, *d* = 0.73). Error rates were also slightly higher on neutral-prime trials than on prime-compatible trials, but this difference was not significant (*t*(29) = -1.7, *p* = 0.1, *d* = 0.28).

For the subjective difficulty ratings we also observed a main effect of prime-response compatibility (*F*(2, 28) = 9.60, *p* < .001, ηp^2^ = .407). Due to the circular nature of the scale, ratings lie between 0 (easy) and 360 degrees (difficult). Participants rated prime-incompatible trials (*M* = 156.6) as significantly more difficult than prime-compatible trials (*M* = 146.3; *t*(29) = 4.1, *p* < 0.001, *d* =0.20) and more difficult than neutral trials (*M* = 145.8; *t*(29) = -4.3, *p* < 0.001, *d* = 0.21). Ratings for neutral-prime trials did not differ from ratings for prime-compatible trials (*t*(29) = -.29, *p* = 0.77, *d* = 0.01).

#### 3.1.2 Prime visibility

Based on the data of the prime visibility task, a *d’* value was computed for each participant as an index of prime visibility. The *d’* values were not significantly different from chance level performance (i.e., zero; mean *d’* = 0.077, *sd* = 0.37; one-sample t-test, *t*(29) = 1.13, *p* = 0.27). Thus, it can be concluded that participants show no reliable sign of awareness of the direction of the prime stimuli. Furthermore, when correlating the compatibility effect in the subjective ratings with the individual *d’* values, we found no significant correlation (*r*(28) = .12, *p* = .54), indicating that the subjective ratings were not influenced by prime visibility.

### 3.2 fMRI

#### 3.2.1 Region of Interest (ROI) Analysis Results

In our previous study, which was identical to the current work except that we did not query subjective difficulty, we observed that the rostro cingulate zone (RCZ) and the anterior insula (AI) both were sensitive to conflicts in response selection (Teuchies et al., 2016). Therefore, in a first set of analyses, we focused specifically on these two brain regions. To do so, we extracted single-trial beta estimates from the RCZ (MNI: 6 20 43) and the AI (MNI: -36 20 -1), both defined a-priori based on our previous study. We then used linear mixed models to examine whether these regions are sensitive to differences in subjective difficulty.

An analysis predicting activity in RCZ by difficulty ratings showed a significant effect of difficulty ratings, *F*(1,28.23) = 17.71, *p* < .001. As can be seen in Figure 2A, the easier a trial was judged to be, the lower the activity in RCZ. A similar analysis predicting activity in the anterior insula by difficulty ratings also showed a significant effect, *F*(1,28.29) = 35.87, *p* < .001. As can be seen in Figure 2B, increased subjective difficulty (i.e., lower ratings) was associated with enhanced activity in anterior insula.

**Figure 2.**
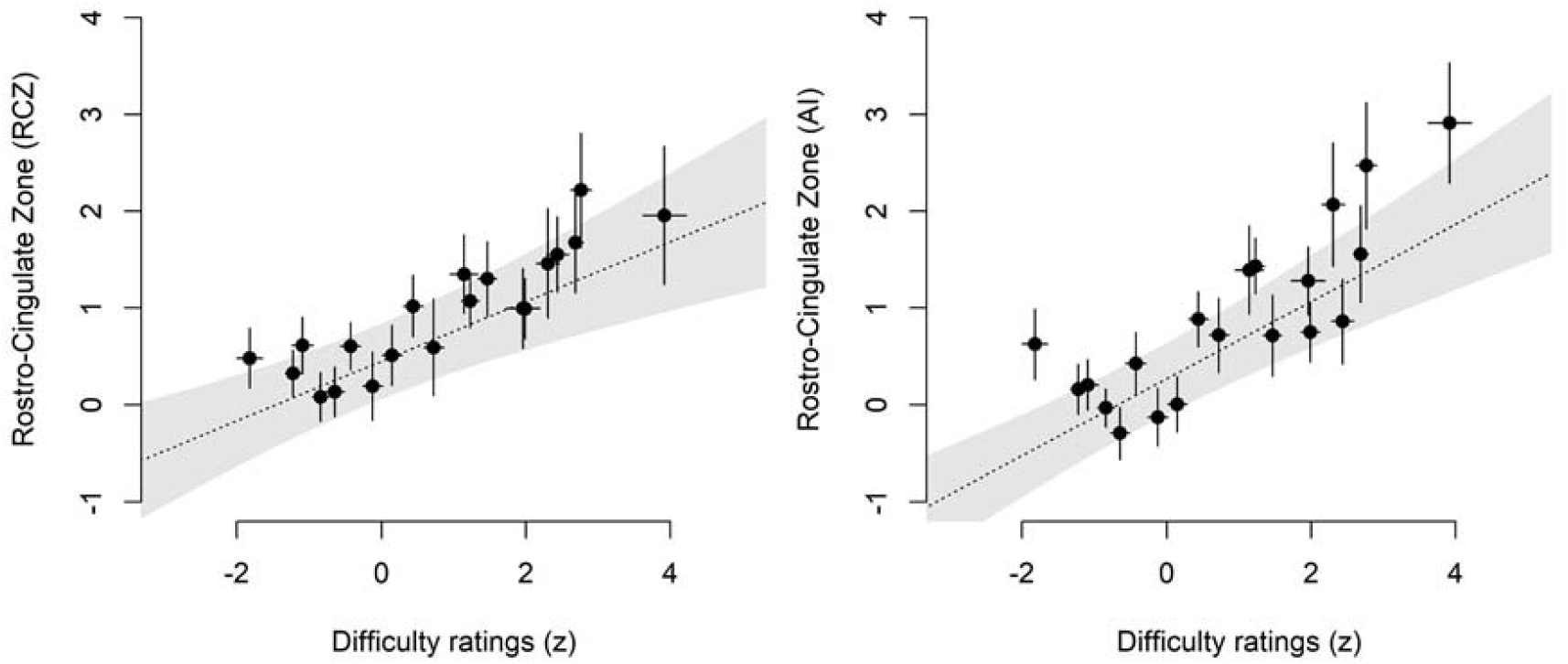
Relation between subjective difficulty ratings and activity in rostro-cingulate zone (A) and anterior insula (B). Dots show the average neural activity in the specified ROI as a function of average difficulty rating, divided in twenty equal-sized bins; dotted lines show the fixed effect slope from the mixed model fit; errors bars and shades reflect standard errors.

Although this analysis is an important first step, it cannot unravel whether activity in RCZ and anterior insula reflects actual variation in difficulty ratings, or whether both are driven by another variable that could in principle covariate with difficulty (such as prime-response compatibility or reaction time; Desender, Van Opstal, & Van den Bussche, 2017). Therefore, in a second set of analyses, we looked whether both these brain regions showed activity *uniquely* correlated with subjective ratings, that is, after controlling for the variables compatibility and reaction times. To do so, we extended the mixed regression models reported before and included compatibility and reaction times.

First, we report the results of a model predicting single-trial RCZ activity by compatibility (3 levels: congruent, neutral or incongruent), reaction times, and subjective difficulty ratings. Replicating our previous work, there was a main effect of compatibility, *F*(2,5814.2) = 4.508, *p* = .011. This main effect reflected that RCZ activity was higher on incongruent trials than on congruent trials, *z* = 2.45, *p* = .014, and on neutral trials, *z* = 2.75, *p* = .006, whereas congruent and neutral trials did not differ, *p* > .757. We also observed a significant main effect of reaction times, *F*(1,33.4) = 27.02, *p* < .001, reflecting increased RCZ activity for trials with slower reaction times. Critically, even after controlling for both these factors we still observed a significant main effect of difficulty ratings, *F*(1,27.9) = 9.80, *p* = .004. The interaction between compatibility and difficulty ratings did not reach significance, *p* =.072.

Second, highly similar results were found in a model predicting single trial anterior insula activity by compatibility (3 levels: congruent, neutral or incongruent), reaction times and difficulty ratings. Also here, we observed a main effect of compatibility, *F*(2,5825.7) = 5.11, *p* = .006, reflecting that activity in anterior insula was higher on incongruent trials than on congruent, *z* = 2.13, *p* = .033, and neutral trials, *z* = 3.14, *p* = .001, whereas congruent and neutral trials did not differ, *p* > .302. We also observed a significant main effect of reaction times, *F*(1,5654.4) = 43.97, *p* < .001, reflecting increased AI activity for trials with slower reaction times. Critically, even after controlling for both these factors we still observed a significant main effect of difficulty ratings, *F*(1,31.3) = 18.91, *p* <.001. The interaction between difficulty ratings and congruency did not reach significance, *p* = .085.

#### 3.2.2 Whole-Brain Analysis Results

In the ROI analyses, we found that difficulty ratings significantly predicted activity in RCZ and anterior insula even when controlling for RT and compatibility. To corroborate these findings, we next performed two whole-brain univariate analyses. We first looked for brain regions where activation magnitude was correlated with subjective ratings. This analysis revealed a large set of regions with significant activation, including the RCZ and insula (see Figure 3). When using a more conservative threshold, clusters surviving correction were located in the RCZ [MNI 3 23 55], right insula [MNI 51 17 4] and left insula [MNI -36 23 -2]. This first whole-brain analysis corroborates the previous analyses showing that increased activity in the RCZ and insula is related to increased subjective difficulty.

**Figure 3.**
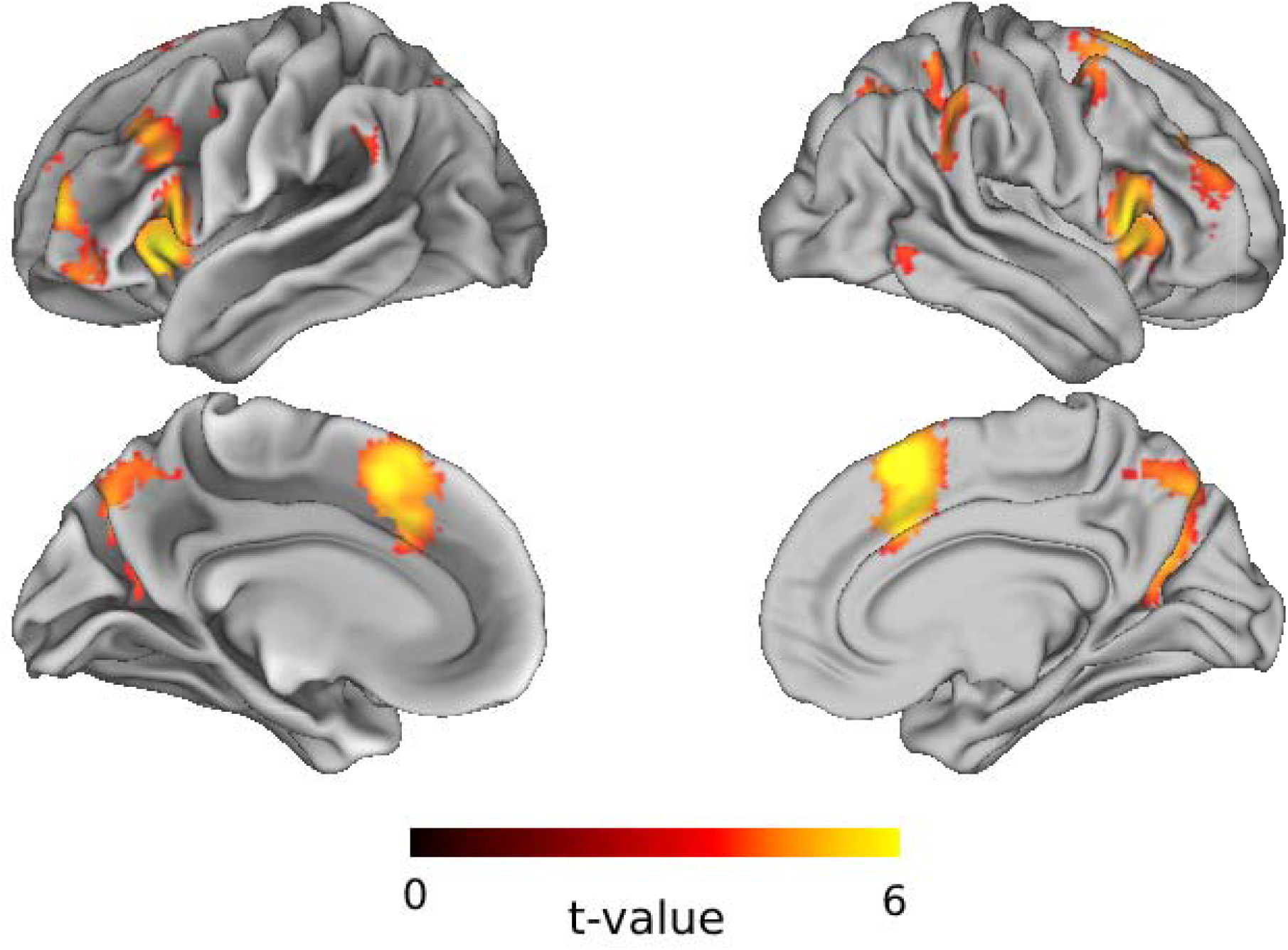
GLM contrast for the effect of subjective difficulty. Warm colors show regions where activation magnitude is correlated with difficulty ratings (primary voxel threshold [p < 0.001 uncorrected] and cluster-defining threshold [FWE p < .05]).

Similar to the ROI analyses, we then looked for brain regions that showed activity uniquely correlated with subjective ratings. To do so, we performed a GLM analysis with one regressor for each compatibility level (compatible, incompatible, neutral) and two parametric modulators: 1) reaction time and 2) subjective ratings. Importantly, the order of the parametric modulators matters, as the second one will only operate over the variance not explained by the first modulator. Therefore, by looking at activity correlated with subjective ratings, we were able to ascertain what brain regions code for subjective ratings, while controlling for reaction times and compatibility. Results from this model showed that the parameter of subjective rating residuals revealed again a set of frontoparietal regions (see Figure 4), which included significant clusters of FWE corrected activation in the RCZ [MNI -3 14 52] and the left [MNI -33 26 5] and right anterior insula [MNI 57 23 7]. The left anterior insula cluster is closely located to the anterior insula [MNI -36 20 -2] that we observed in our previous study (Teuchies et al., 2016). These results indicate that the RCZ and the anterior insula showed increased activity with increased subjective difficulty, independent from prime-response compatibility or reaction times.

**Figure 4.**
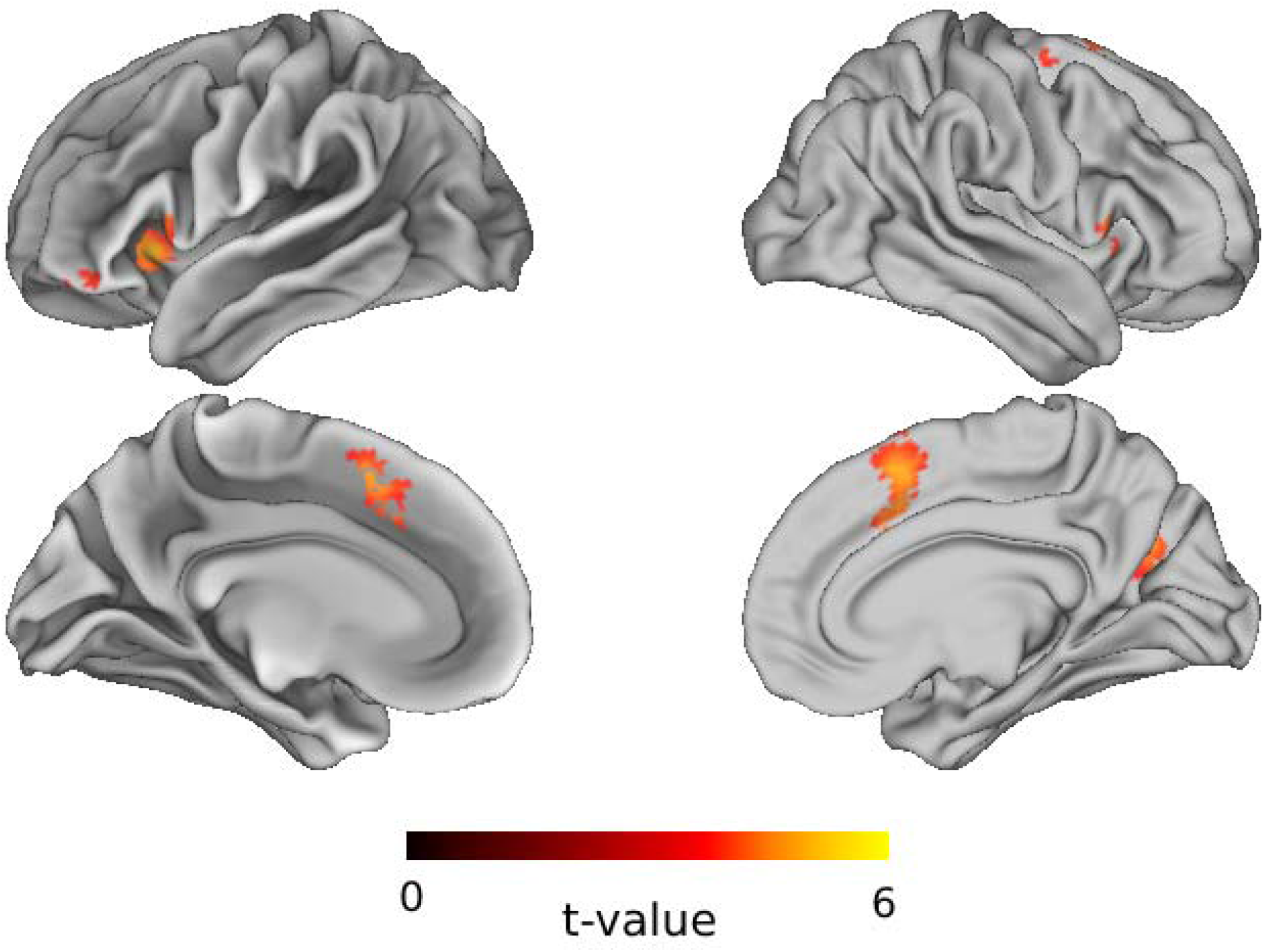
Main areas of interest showing higher activation when the sense of subjective difficulty increased, independent from prime-response compatibility and after regressing out the effect of reaction times. Warm colors show regions where activation magnitude is correlated with difficulty ratings (primary voxel threshold [p < 0.001 uncorrected] and cluster-defining threshold [FWE p < .05]).

#### 3.2.3. Mediation analysis

To shed light on the directionality between both identified brain regions, we next performed causal mediation analysis. First, we tested the hypothesis that the influence of the RCZ on ratings is mediated by the anterior insula. A prerequisite for mediation analysis is that all three paths are significant. Indeed, in the mediator model, the RCZ predicted activity in the anterior insula, *F*(1,28.4) = 511.08, *p* < .001, and in the outcome model subjective ratings were significantly predicted by the anterior insula, *F*(1,5813) = 11.05, *p* < .001, and RCZ, *F*(1,5814.1) = 3.011, *p* = .083. Results of the mediation analyses showed that, conditional on compatibility and RTs, there was a significant part of the influence of the RCZ on subjective ratings that was mediated by activity in the AI, β = .451, *p* < .001, whereas there was no direct effect of the RCZ on subjective rating, β = .404, *p* = .078. These results are shown in Figure 5. Because mediation analysis is a correlational technique, we also tested the reversed causal flow, namely that the influence of the AI on ratings is mediated by the RCZ. This analysis showed a highly significant direct effect of AI on ratings, β = .701, *p* < .001, and no mediation effect by the RCZ, β = .209, *p* = .099. In sum, results from the mediation analyses suggest that the influence of the RCZ on ratings is mediated by activity in the AI.

**Figure 5.**
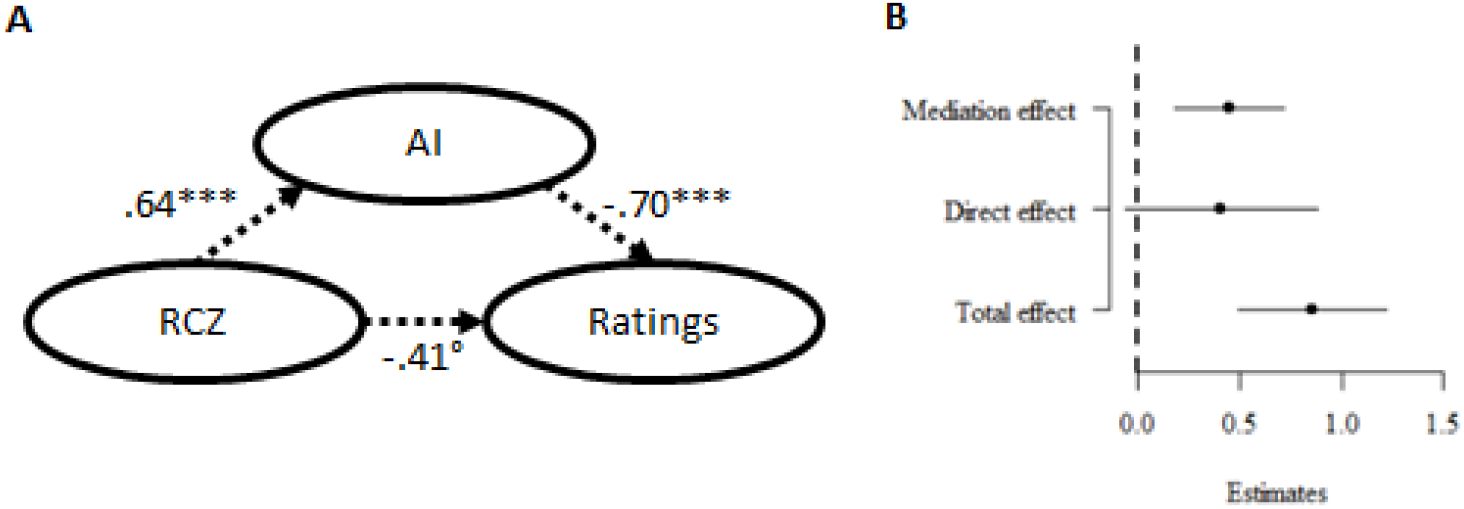
Results of the mediation analysis testing whether the influence of RCZ on ratings is mediated by AI. Unstandardized regression coefficients are shown from the mediation and the outcome mixed regression models. Degrees of freedom for calculating significance were based on Satterthwaite’s approximation. Effect sizes from the causal mediation analysis are shown in (b). Error bars reflect quasi-Bayesian 95% confidence intervals. Note: ***p<.001, °p<.1

## 4. Discussion

In the current study, we set out to test which brain regions are involved in self-monitoring difficulty in action selection. To accomplish this, a subliminal response priming task was used that influences the difficulty of action selection. The benefit of this task is that participants remained unaware of the conflict-inducing stimulus itself, thereby eliminating the possibility that participants rated their sense of subjective difficulty based on perceiving the conflict-inducing stimuli or based on how they believe they should respond. In the current study we observed that participants reported increased subjective difficulty on trials in which subliminal response conflict was induced. In a first analysis step, activation in the RCZ was found that was related to the raw subjective difficulty ratings. In a second step, we then wanted to unravel activity specific to subjective ratings, which was not explained by prime-response compatibility or reaction times. This analysis showed that both the RCZ and the anterior insula were related to unique variation in subjective ratings. Further, replicating previous work, we found that activation in both these regions was increased in the presence of subliminal response conflict. In the remainder of this section, we discuss how these findings are compatible with a grounding of metacognitive experiences of difficulty within interoceptive signals.

### 4.1. Metacognitive computations of subjective difficulty

The RCZ is part of the anterior cingulate cortex, which is a central hub in the cerebral cortex. This brain region plays a key role in cognitive control processes, as it is argued to be involved in conflict processing (Botvinick et al., 2001), in computing the expected value of control (Shenhav et al., 2013), in detecting violations from predictions (Alexander & Brown, 2011), and in effort processing (Walton et al., 2002). These different functions have been integrated by positing that the ACC controls the degree of effort invested in a certain task (Holroyd & McClure, 2015). Indeed, focal damage to rat medial frontal cortex decreased the frequency of high effort responses to obtain a reward. Within this framework, the sensitivity of the anterior cingulate to response conflict results from an increased need for effort in difficult trials (i.e., conflict trials). Confirming this prediction and replicating previous work, the current study observed higher activation in the RCZ for incompatible trials compared to compatible and neutral trials. Given that subjective difficulty judgments track response conflict, a relation between subjective difficulty and RCZ was expected. Critically, however, we were able to demonstrate that the relation between difficulty ratings and RCZ was present, even after controlling for the influence of prime-response compatibility and reaction times. This shows that metacognitive computations of subjective difficulty do not merely track experimental manipulations, rather, they are based on brain regions, such as the RCZ, that code for the required degree of effort, over and above that induced by the experimental manipulation.

Whether or not metacognitive computations of subjective difficulty are directly related to RCZ activity or only indirectly remains an open question. One interesting possibility is that the RCZ only codes for the required degree of effort, and this is subsequently implemented by other brain regions. In this regard, it is interesting that apart from the RCZ we also observed that the anterior insula was sensitive to subjective difficulty ratings, over and above the effect of response conflict and reaction times. Note that the anterior insula was not observed in the analysis where we only looked at subjective ratings (i.e., uncontrolled for compatibility and RTs). One explanation for this might be that by factoring out variance from RTs and compatibility from subjective ratings, the remaining variance reflects a more veridical measure of subjective difficulty (e.g., uncontaminated by RTs) and therefore we have more power to detect effects. The anterior insula is a key brain region involved in interoceptive awareness (Craig, 2009; Grupe & Nitschke, 2013; Gu, Hof, Friston, & Fan, 2013). Interoception can be described as the sense of the physiological condition of the body, or the perception of sensory events occurring within one’s body (Craig, 2002; 2003; Grupe & Nitschke, 2013). The anterior insula is thought to monitor and control internal, embodied states, such as the degree of arousal. Thus, when the RCZ detects the need for increased effort allocation, this might subsequently be implemented by the anterior insula that increases arousal, via interactions with the sympathetic nervous system. This interpretation is further supported by the results of the mediation analysis carried out in the current study, which suggest that the influence of the RCZ on subjective difficulty ratings is mediated by activity in the AI. Given that humans are typically unaware of their own brain activity (Prinz, 1992), this raises the interesting possibility that metacognitive evaluations of difficulty are based on bodily signals in response to required effort. Thus, when judging whether a trial was easy or difficult, participants might integrate, among other things, their autonomous bodily reactions towards subliminal response conflict (i.e. cardiac acceleration, increased skin conductance; Allen et al., 2016; Hauser et al., 2017) in order to come to a single judgment of difficulty. Indeed, a recent study demonstrated that participants relied on motor activity in their response effectors (i.e., in the thumbs of both hands) when judging the difficulty of a trial (Questienne et al., 2017).

### 4.2. Domain-general versus domain-specific metacognition

In recent years, the metacognitive evaluation of performance has been tackled from different angles. This has raised the question whether this metacognitive evaluation of performance is supported by a set of domain-general mechanisms, or whether there is domain-specificity. To tackle this question, McCurdy and colleagues (2013) compared metacognition about visual performance with metacognition about memory performance. Although metacognitive performance was correlated across both domains (see also Faivre et al., 2017; Song et al., 2011), different neural structures were involved in each. Whereas metacognitive performance about visual decisions was related to volume in anterior prefrontal cortex (see also Fleming et al., 2010), metacognitive performance about memory decisions was related to the precuneus. In a subsequent study, metacognitive performance about visual decisions was linked to white matter microstructure in the ACC whereas metacognitive performance about memory was linked to white matter microstructure in the inferior parietal lobule (Baird et al., 2014). The current work adds to this debate, by demonstrating that in a different type of self-evaluation, subjective difficulty in decision making, both RCZ and anterior insula are involved. The latter is particularly interesting, because although the anterior insula is critically involved in self-referential processes such as self-awareness (Craig, 2009), previous studies did not, to our knowledge, implicate the anterior insula in the metacognitive evaluation of performance. As such, the current findings lend further support for the domain-specific view of metacognition.

## 5. Conclusion

In the current work, we observed that the subjective sense of difficulty is represented in the RCZ and the anterior insula, two regions that are functionally closely connected. Importantly, this was observed when controlling for prime-response compatibility and reaction times. Since the RCZ and the anterior insula typically activate in unison, future research is needed to test our hypothesis that the RCZ codes for the required level of effort and the anterior insula implements this by increasing arousal, which is what participants become aware of.

## 6. Acknowledgements

The research reported in this paper was funded by the Interuniversity Attraction Poles Program initiated by the Belgian Science Policy Office (IUAPVII/33). K.D. is an FWO [PEGASUS]^2^ Marie Skłodowska-Curie fellow (grant number 12T9717N). C.G.-G. was supported by the European Union’s Horizon 2020 research and innovation programme under the Marie Sklodowska-Curie grant agreement no. 835767. MB is supported by an Einstein Strategic Professorship of the Einstein Foundation Berlin.

